# Symmetry breaking and social coordination in children: Using group theory to understand emergent patterns of multi-agent coordination

**DOI:** 10.64898/2026.08.11.743695

**Authors:** Akifumi Kijima, Motoki Okumura, Hiroyuki Shima, Rachel W. Kallen, Michael J. Richardson, Yuji Yamamoto

## Abstract

Predicting patterns of behavioural coordination that emerge in small interacting groups is challenging because goal-directed social action is shaped by complex reciprocal and compensatory dynamics. In this study, we examined whether formal symmetry principles derived from group theory could explain coordination patterns among children performing a triadic jumping task. We investigated how geometric symmetries of the task environment and dispositional (a)symmetries associated with leader-follower tendencies jointly constrain collective behaviour. Forty-seven children were classified into symmetric or asymmetric triads based on teacher evaluations of leadership dispositions. Each triad completed multiple trials of a synchronized jumping game requiring movement between adjacent hoops arranged in triangular or square configurations. Results showed that temporal asymmetries in inter-child movement (first, second, or last to jump) were consistent with group-theoretic predictions. In triangular configurations, observed asymmetries corresponded to the highest-order subgroup defined by task and dispositional symmetries. These findings demonstrate that environmental symmetry exerts a hierarchically dominant constraint on collective coordination, within which actor dispositional (a)symmetries further modulate emerging patterns.

## 1. Introduction

Human social movement coordination is a complex process with the stability and patterning of such behaviour not only shaped by the reciprocal interactions that occur between multiple individuals, but by the constraints of the task environment and the dispositional differences of the individuals involved. While a great deal of prior research has sought to understand these complex dynamical processes, this research has primarily focused on dyadic coordination (i.e., synchronization or coordination between two individuals[1, 2, 3, 4, 5, 6, 7]). This is despite the fact that real-world social interaction often involves three or more people, requiring the simultaneous stabilization of multiple factors and relationships. This is true even for triadic coordination tasks, which entail a significant degree of additional complexity compared to dyadic interaction (as does any three-body problem), demanding the dynamic balancing of several physical and informational factors, and individual differences, leading to a diverse range of context dependent, behavioral motifs.

Key to understanding such behavior is adopting a formal approach that both defines key task constraints and individual differences, and predicts the possible patterns these constraints and differences entail. Recently, Richardson and Kallen [8, 9] and others[10, 11, 12, 13] have proposed that the formal principles of symmetry (i.e., group theory) and symmetry breaking provide the ideal framework for understanding and predicting the patterns of behavioural coordination that can emerge in social and multi-agent interactions. Indeed, just as the principles of symmetry and symmetry breaking play a foundational role in understanding complex phenomena in the physical and biological sciences[14, 15, 16, 17, 18, 19], they also appear to provide a foundational way of understanding why and when certain patterns of group behaviour emerge within a given task or social context[9].

The aim of the current work was to explore this possibility by examining the degree to which the patterns of behavioural coordination that emerge during small group activity can be understood using the theoretical and formal principles of symmetry and symmetry breaking. More specifically, the current study used symmetry groups to explore the degree to which the physical task (a)symmetries of a three-person (triadic) jumping game in combination with the psycho-social (a)symmetries that characterize differences in the leader-follower tendencies of the children completing the game define the specific patterns of coordination that emerge.

### Triadic jumping task

As an initial investigation into whether symmetry principles and group theory can be used to understand the patterns of behavioral coordination that emerge between small groups of interacting individuals, we previously conducted an experiment examining whether the geometric symmetry of a triadic jumping task could predict the leader-follower relationship among adult jumpers when they successfully completed the task.

In this experiment, we arranged four distinct jumping “workspaces”, each with a different geometrical configuration, including the triangular and square setups shown in Fig. 1. The adults in each triad were instructed to repeatedly exchange positions by jumping in synchrony in response to a metronome cue presented every 3 seconds, as illustrated in Fig. 1(A). Importantly, because participants were instructed not to communicate, the group could successfully complete the task only if all three individuals spontaneously converged on the same movement direction. This required one participant to initiate the jump slightly before the others, thereby indicating the direction that the group should follow. Otherwise, participants collided, failed the trial, and were required to restart it (see Fig. 1(A), “Triadic jumping task” in “Method Details” for further details).

**Figure 1:**
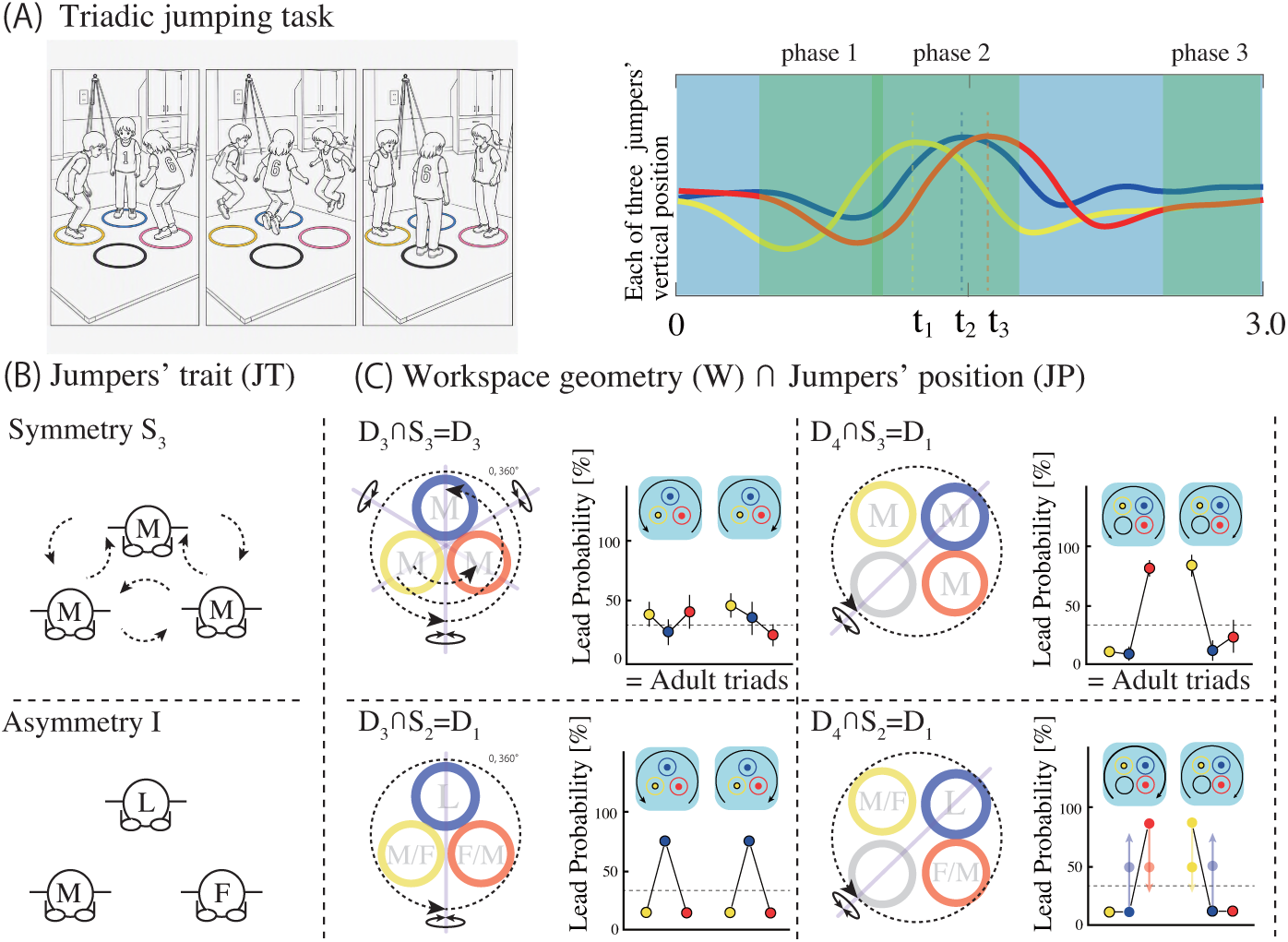
An illustration of the Triadic jumping task and hypothesis. **(A)** Time series of vertical body position in one cycle of triadic jumping: Phase 1, arm swing and knee flexion; Phase 2, center of mass (CoM) peak; Phase 3, CoM returns on landing. **(B)** The symmetry group (*S*) of the jumpers’ trait for Symmetric and Asymmetric triads. L: classroom leaders, M: moderates and F: followers. Symmetric triads can be represented by *S*_3_, which has order 6, while asymmetric triads can be represented by *I*, which has order **1. (C)** Hypothesis (line plot) on the lead probability among three jumpers performing triadic jumping in triads with either Symmetrical or Asymmetrical leader-follower traits under two geometric configurations. We hypothesize that the orderly leader-follower pattern among the jumpers corresponds to the highest order common subgroup within the dihedral symmetry groups (*D*) of the workspace geometry and the symmetry group (*S*) of the jumpers’ traits. For asymmetric triads, the positions of moderates and followers are swapped with equal probability. Therefore, we predicted an orderly leader-follower pattern based on the Jumpers’ position symmetry (JP) as *S*_2_, which differs from the Jumpers’ trait symmetry (JT = *I*). See Appendix A for further details.

The findings indicated that the lead jumper (i.e., the individual who jumped earlier than the other two) was fully determined by the symmetry of the workspace geometry (see, upper two panels in Fig. 1(C)). More specifically, the position of the lead jumper was consistent with the pattern predicted by the isotropy subgroup with the highest level of symmetry, determined from the discrete symmetry groups of the workspace geometry and the functional dynamics of the participants.

With regard to the latter, the three adult jumpers were randomly grouped healthy adults who, as jumpers (JT in the upper panel of Fig. 1(B) and the first and second rows, merged under the label *Adult & Symmetric Children* in the first column of Table 1), could be considered fully symmetric. That is, the functional symmetry of the three jumpers could be represented by the discrete symmetry group *S*_3_.

**Table 1:**
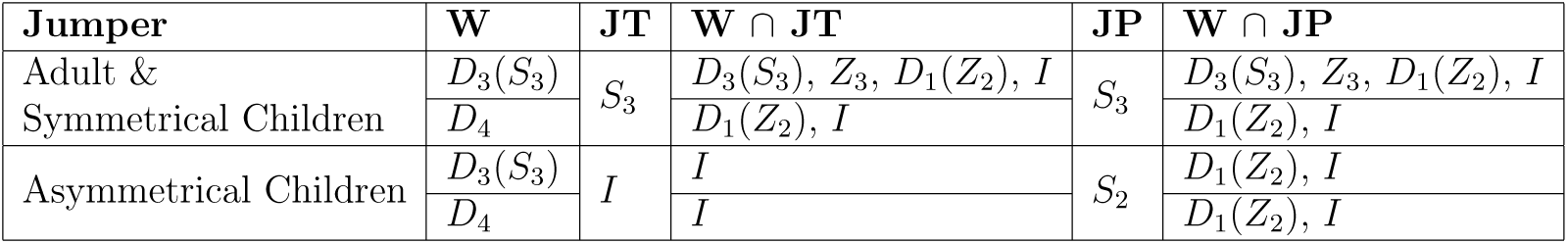
Possible symmetry (Group) predictions for the triadic jumping task with different leader-follower trait symmetries. W: Workspace symmetry, JT: Jumpers’ trait symmetry W*∩*JT: Isotropy subgroup between workspace and jumpers’ trait symmetry. JP: Jumpers’ position symmetry, W*∩*JP: Isotropy subgroup between workspace and jumpers’ position symmetry.

| **Jumper** | **W** | **JT** | **$W \cap JT$** | **JP** | **$W \cap JP$** |
| --- | --- | --- | --- | --- | --- |
| Adult &<br>Symmetrical Children | $D_3(S_3)$ | $S_3$ | $D_3(S_3), Z_3, D_1(Z_2), I$ | $S_3$ | $D_3(S_3), Z_3, D_1(Z_2), I$ |
| | $D_4$ | | $D_1(Z_2), I$ | | $D_1(Z_2), I$ |
| Asymmetrical Children | $D_3(S_3)$ | $I$ | $I$ | $S_2$ | $D_1(Z_2), I$ |
| | $D_4$ | | $I$ | | $D_1(Z_2), I$ |

With regard to workspace configuration, for the triangle workspace, as shown in the left two panels of Fig. 1(C), the geometrical symmetry of this workspace was defined by the discrete symmetry group *D*_3_. When considered with respect to the symmetry of the jumpers (*S*_3_), as shown in the upper panel of Fig. 1(B), the corresponding isotropy subgroups that defined the triadic jumping task in the triangle workspace were therefore *D*_3_(*S*_3_), *Z*_3_, *D*_1_(*Z*_2_), and *I*, as indicated in W*∩*JT and W*∩*JP, Table 1. Translated into behavioural predictions about who would jump first, this predicted that each individual in the group was equally likely to jump first on any given jump (metronome cue), which is exactly what was observed — the permutation symmetry of the group (i.e., *S*_3_=*D*_3_) — was preserved over a jumping sequence, as indicated in the left upper panel of Fig. 1(C).

In contrast, although the geometrical symmetry of the square workspace (four-hoop: right two panels of Fig. 1(C)) was defined by the discrete symmetry group *D*_4_, the isotropy subgroups of this symmetry group in relation to the symmetry of the three jumpers (*S*_3_: upper panel of Fig. 1(B)), were only *D*_1_(*Z*_2_) and *I* as indicated in the second row in W*∩*JT and W*∩*JP, Table 1 (also see, right upper panel of Fig. 1(C)). That is, the highest-order subgroups for the square workspace was *D*_1_(*Z*_2_), which corresponded to only two individuals being permutable i.e., jumpers in the red and yellow hoop in right upper panel of Fig. 1(C). Consistent with this lower-order symmetry prediction, see plots in right upper panel of Fig. 1(C), one jumper consistently acted as the leader, while the other two were rarely ever the lead jumper. That is, the explicit symmetry break the 4-hoop + 3-jumper condition resulted in one of the individuals next to the open hoop emerging as the leader, and continuing to act as the leader thereby ensuring task success across jumps and trials.

### Current Study: Effect of “asymmetric” leader-follower individuals on group coordination

The aim of the present study was to validate and extend our previous work by exploring whether individual differences in leader-follower tendencies act as an explicit symmetry-breaking factor that shapes the coordination patterns emerging during the triadic jumping task, and whether these coordination patterns can be understood using symmetry groups. To address these questions, we recruited elementary school children and assigned them to triads based on their leader-follower tendencies in daily classroom activities. Each child was classified by their classroom teacher as a leader, a moderate, or a follower. Children were chosen because their dispositional tendencies are more naturally expressed, and because in the early elementary school years they still rely largely on external norms and expectations. In this developmental context, these leader-follower relationships co-evolve through mutual challenge and support, forming a foundation for social growth in both leadership and followership [20]. At this stage, dispositional differences are relatively large but, as children encounter varied social and school environments, these differences tend to average out, providing a foundation for the development of more self-determined values and behaviors [21, 22, 23, 24].

Importantly, the differences in leader-follower trait symmetry (JT) can be represented by the identity group *I*, as shown in JT in the third and fourth rows in Table 1 which are merged under the label *Asymmetric Children* and the lower panel of Fig. 1(B), meaning that each individual has a unique individual identity/disposition. This is in contrast to groups of children who did not demonstrate any clear leadership tendencies and could be approximately represented by *S*_3_; just as the adult groups in [12]. Of particular interest, here, was when differences in leader-follower dispositions (i.e., asymmetric triads) would operate to break the symmetry of the group and, thus, lead to coordination patterns defined by *I* in W*∩*JT. Note that this represents the isotropy subgroup of each workspace’s geometric symmetry (*D*_3_ and *D*_4_) combined with *I* (leader *→* moderate *→* follower). In other words, by confirming the actual temporal pattern of triadic jumping matching W*∩*JT in the third and fourth columns in Table 1 and depicted in the lower panel of Fig. 1 (B) we aimed to test whether the constraints imposed by the participants’dispositional traits determined the observed coordination pattern of the triadic jumping.

To compare the influence of workspace geometric symmetry with the independent effects of the jumpers’ leader-follower dispositions, we counterbalanced the arrangement of moderates and followers. Specifically, moderates and followers were placed in the red and yellow positions shown in the two lower panels of Fig. 1(C) with equal probability, while classroom leaders were always positioned in the blue hoop. In the triangle workspace, there were no open spaces; this resulted in all three positions being perfectly symmetrical with each other. In contrast, in the square workspace, only the red and yellow positions (hoops where two jumpers with lower leadership tendencies were located) were adjacent to open spaces, while the blue position had no open space on either side (a hoop where the jumper with the strongest leadership tendency was located). Thus, the impact of individual dispositional traits should be different across conditions.

For the triangular condition, where the symmetry is conserved with respect to the hoop configuration, children in symmetric groups should be equally likely to adopt the role of leader, as observed in [12]. Thus, *D*_3_(*S*_3_) should be observed across jumps and trials, with the identity of the first jumper reflecting *S*_3_ symmetry from jump to jump and trial to trial. For the asymmetric groups, however, the difference in leader-follower disposition should explicitly break the symmetry, resulting in the child judged to have the strongest leadership tendencies predominantly jumping first. That is, *D*_1_(*Z*_2_) should be observed if jumpers’ position symmetry (JP: *S*_2_) constrains jumping order. On the other hand, *I* should be observed if jumpers’ trait (JT) independently constrains jumping order. Details of the JT and JP symmetries and (highest order) common subgroups with the workspace geometry (W) are provided in Appendix A.

In contrast, for the square configuration, the symmetry is explicitly broken by the geometric arrangement, such that for symmetric triads, *D*_4_ *∩ S*_3_ = *D*_1_(*Z*_2_) should be observed (upper right panel of Fig. 1(C) and W*∩*JP in the 2nd row of Table 1). This pattern of results was also possible for asymmetric triads because the classroom leader was always positioned in the hoop diagonally opposite the open space, while the positions of the moderates and followers were counterbalanced. Consequently, the jumper-position symmetry (JP) remained *S*_2_, as detailed in Appendix A. Observing this result would imply that the coordination pattern that emerged was unaffected by the leader-follower asymmetries and that the coordination patterns that emerged were a function of W*∩*JP in the 4th row of Table 1. Importantly, however, if asymmetries in the jumpers’ traits do influence the resulting coordination pattern, a pattern consistent with I should be observed (W*∩*JT in the fourth row of Table 1).

## 2. Results

To determine whether the observed patterns of triadic coordination that emerged among the children were consistent with the possible symmetry group predictions outlined above and in Table 1 and Figure 1, we analyzed the data in three ways. We first analyzed asymmetric triads to examine how children’s dispositional traits, rather than workspace symmetry, influenced the probability of initiating a jump. Of particular interest here was the degree to which the jumping order of classroom leaders, moderates and followers was consistent with the patterns defined in Table 1. We then examined the probability that each child would jump first as a function of trait symmetry (symmetric vs asymmetric triads) and jumper hoop position for both the triangle (*D*_3_) and square (*D*_4_) workspaces. This analysis was done in order to identify whether the role of jumper position in the pattern of coordination observed and, moreover, the degree to which asymmetries in jumper trait superseded (a)symmetries in jumper position (particularly for the square workspace). Finally, we examined the temporal lead-lag relationships among jumpers to further validate the degree to which asymmetries in jumper trait and/or position defined the functional asymmetries in the coordination dynamics required to achieve task success (i.e., at least one child must jump first). Throughout this study, “who jumped first” refers to the identity of the child who first initiated the jump during a given jump. The corresponding probability therefore represents the proportion of jumps in which each child initiated the jump first.

### 2.1. Effects of trait asymmetries on who jumps first

Figure 2(A) shows the probability of which child jumped first as a function of leader-follower characteristics and workspace geometry for asymmetric groups. A two-factor repeated measures analysis of variance was employed to examine the effects of workspace geometry (Geo-symmetry: triangular and square) and jumper trait (Jumper-trait: Classroom leaders, moderates, followers) on the probability of a jumper being the lead jumper (Mauchly’s sphericity tests were not significant for both measures and the interaction effect).

**Figure 2:**
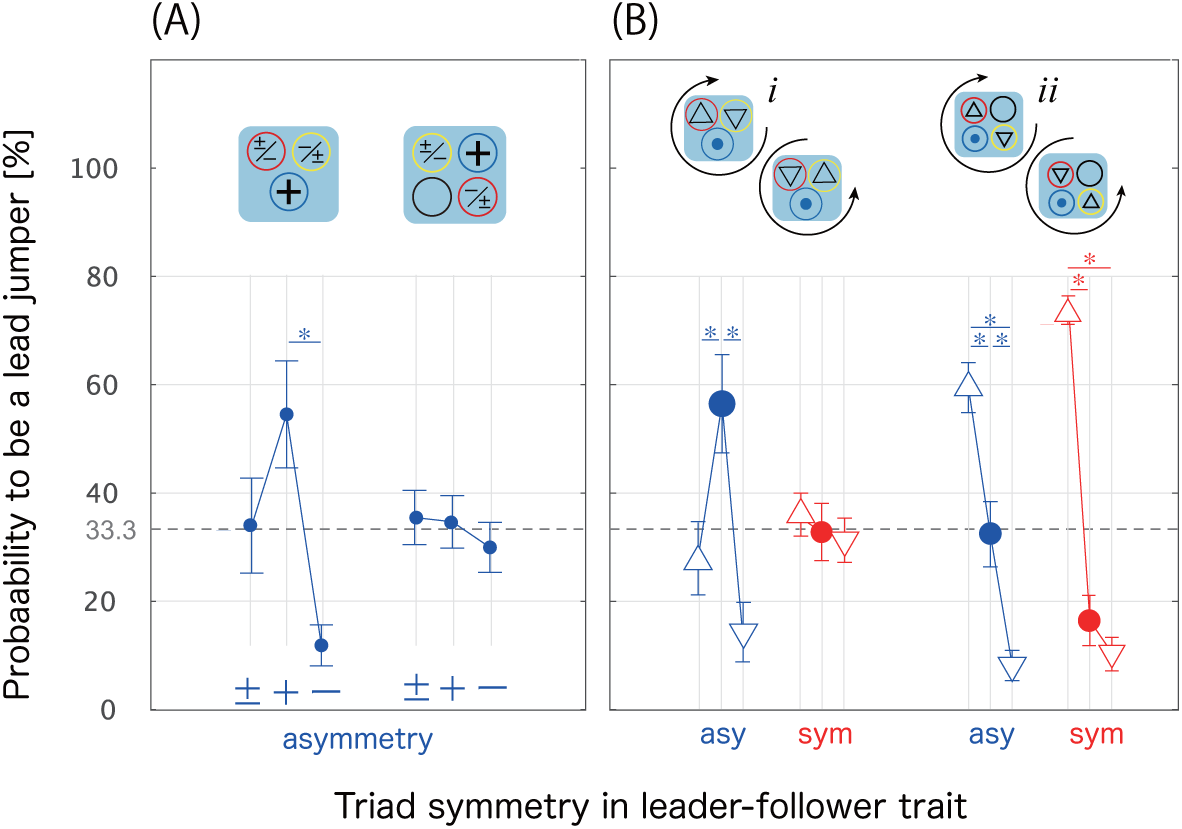
Probability of being the lead jumper in Triangle and Square workspace, with * indicating a significantly higher probability between jumpers. **(A)** *Eflects of the Jumpertrait* for asymmetric triads in the Triangle (left) and square workspaces (right), with leaders denoted by ‘+’, moderates by ‘*±*’ and the followers by ‘*−*’. For the Triangle workspace, classroom leaders exhibited a significantly higher probability of jumping first compared to followers. However, there was no significant difference observed between leaders and moderates. For the square workspace there was no significant difference in who jump first as a function of jumper trait. **(B)** *Effects of the geometric, Jumper Position symmetry* for asymmetric and symmetric trials for the Triangle (i) and Square (ii) workspaces. For the Triangle workspace, each child was equally likely to jump first for symmetric groups, whereas the classroom leaders exhibited a significantly higher probability of jumping first compared to moderates and followers in asymmetric groups. There was no significant difference observed between the moderates and followers who were always located *in front of* and *behind of* the classroom leaders with respect to the direction the triad jumped. For the Square workspace, significant differences were observed between all three jumpers in Asymmetric triads. However, for Symmetric triads, only jumpers assigned in one hoop positioned *in front of* an open hoop (red for clockwise jumps and yellow for counterclockwise jumps) demonstrated a significantly higher probability of jumping first compared to jumpers assigned in the other two hoops. Note: In the Triangle and Square workspaces, the moderates and followers were counterbalanced so that they were placed in the red and yellow hoops with equal probability, respectively. The classroom leader, on the other hand, was always allocated to the blue hoop with both sides occupied by other two jumpers.

The analysis revealed no main effect of Geo-symmetry (F(1,10)=0, p=1.000, 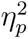=0, 1-*β*=0.050) nor a main effect of Jumper-trait (F(2,20)=2.654, p=0.095, 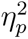=0.210, 1-*β*=0.826), but a significant Geo-symmetry x Jump-trait interaction (F(2, 20)=5.419, p =0.013, 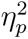=0.351, 1-*β*=0.948). A simple effects analysis revealed that the effect of Jumper-trait was significant only for triangle workspace (F(2,20)=4.105, p=0.040, 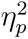=0.291), with Bonferroni post-hoc analysis (*α*=0.05, two-tailed test) indicating that the probability of classroom leaders (54.4%) jumping first was significantly higher than the probability of followers (11.7%), but not significantly higher than that of moderates (33.9%). That is, the jumper-trait asymmetry operated to break the symmetry of the group, with the resulting pattern best captured by *D*_1_(*Z*_2_) or *I* in W*∩*JT in Table 1.

In contrast, there was no difference in the probability of who jumped first for the square workspace (F(2,20)=0.112, p=0.895, 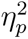=0.011), with each child equally likely to jump first on any given trial. At first glance, this latter result appears to be inconsistent with the predictions in Table 1, highlighting the potential balance (competing influences) that trait asymmetry and geo-symmetry played in structuring the groups jumping dynamics. More specifically, classroom leaders jumped first as often as moderates and followers, despite the geo-symmetry defining them as predicted to be least likely to jump first. However, as we detail below, this result was in part an artifact of the covariance between jumper position and direction, with the probability of who jumped first for asymmetric triads in the square condition best captured by *I* as shown in blue data points in Fig. 2(B)-ii.

### 2.2. Effects of jumper position and trait symmetry on who jumps first

#### 2.2.1. Triangle workspace

Fig. 2(B)-i depicts the probability of who jumped first as a function of jumper position for the triangle workspace. Note that for the asymmetric triads, the blue dot indicates where the children classified as a “classroom leader” were positioned. A two-factor, mixed analysis of variance was employed to examine the effects of Triad-symmetry (asymmetric vs symmetric) and relative position (Jump-position) With regard to observed jump direction on the probability of being the lead jumper, with Jump-position as the repeated measure. Mauchly’s sphericity test for Jump-position was significant (Mauchly’s W=0.616, p=0.009), and thus the Greenhouse-Geisser degrees-of-freedom adjustment was employed for effects that included this factor.

The analysis revealed a significant main effect of Jump-position (F(1.445, 28.896)=4.111, p= 0.038, 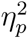=0.171, 1-*β*=0.897) and a significant Triad-symmetry x Jump-position interaction (F(1.445, 28.896)=3.868, p =0.045, 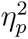=0.162, 1-*β*=0.878). There was no main effect of Triad-symmetry (F(1,20)=1.956, p=0.341, 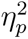=0.046, 1-*β*=0.393). Consistent with the results reported above, a simple main effects analysis revealed that for the triangle workspace the effect of Jump-position was significant for asymmetric triads (F(2,40)=7.8317, p=0.007, 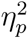=0.281; shown in blue line plot), with Bonferroni post hoc analysis revealing that for the asymmetric triads the child identified as a classroom leader (placed in the blue hoop) had a significantly (*p*=0.017) higher probability (57.1%) of jumping first compared to the other two jumpers. There was no significant difference between the probabilities of the other two jumpers (p*>*0.05). This was true independent of jump direction, with the jumper located in the hoop *in front of* and *behind* the classroom leader with respect to the observed jumping direction jumped first only 28.3% (*△* in Fig.2(B)-i) and 14.6% (*▽* in Fig.2(B)-i) of the time, respectively. As expected, there was no effect of Jump-position for symmetric triads (F(2,40)=0.1472, p=1.000, 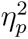=0.007; shown in red line plot), with the jumper in each position equally likely to jump first on any given trial.

Although the observed pattern of coordination for symmetric triads reflected the highest-order isotropy subgroup for both W*∩*JT and W*∩*JP for a JT and JP of *S*_3_, that is, *D*_3_(*S*_3_), the collective results for the triangle workspace better matched the symmetry group predictions in Table 1

W*∩*JT. More specifically, for the asymmetric triads, the observed pattern of results reflected *D*_1_(*Z*_2_), which is the highest order isotropy subgroup for the workspace symmetry *D*_3_(*S*_3_) and the jumper trait symmetry of *S*_2_ in W*∩*JP (the third row in Table 1). This implies that in the presence of a classroom leader, the followers and children with only moderate leader tendencies could be considered equivalent (i.e., per-mutable; recalling that the positions of the moderates and followers were counterbalanced across trial sequence). Moreover, that the presence of a leader in a symmetric workspace acted as a symmetry breaking factor With regard to establishing a functional patterning of the triadic jumping coordination.

#### 2.2.2. Square workspace

Fig. 2(B)-ii depicts the probability of who jumped first as a function of jumper position for the square workspace. Again, a two-factor, mixed design analysis of variance was employed to examine the effects of Triad-symmetry and Jump-position relative to the observed jump direction on the probability of who jumped first. Mauchly’s sphericity test for Jump-position was significant (Mauchly’s W=0.606, p=0.009), and thus the Greenhouse-Geisser degrees-of-freedom adjustment was employed for effects that included this factor.

The analysis revealed a significant main effect of Jump-position (F(1.434, 28.688)=59.973, p= 0, 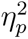=0.750, 1-*β*=1) and a significant Triad-symmetry x Jump-position interaction (F(1.434, 28.688)=3.966, p =0.043, 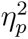=0.166, 1-*β*=0.998). There was no main effect of Triad-symmetry (F(1,20)=2.222, p=0.152, 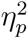=0.100, 1-*β*=0.972).

Despite the significant interaction, a simple effects analysis revealed a significant effect of Jump-position for both asymmetric (F(1.434, 28.688)=22.479, p<0.001, 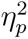=0.529) and symmetric (F(1.434, 28.688)=41.460, p<0.001, 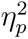=0.675) groups. For asymmetric groups, this reflected the fact that the child positioned next to the open hoop who jumped first defined the direction with which the triads jumped. That is, if the child in the red hoop jumped first the triad tended to jump in the clockwise direction; whereas if the child in the yellow hoop jumped first the triad tended to jump in a counterclockwise direction. Thus, unsurprisingly, Bonferroni post-hoc analysis revealed that the jumper positioned in the hoop *in front of* the open space relative to the direction of the jump, was significantly more likely to jump first (59.5%) compared to the jumpers in the other two positions (p=0.015). Interestingly, the child designated as a classroom leader and who was always positioned in the hoop *diagonal* to the open space, did still jump first significantly more than the jumper in the hoop behind the open space (32.4% vs. 8.1% for *diagonal* and *behind* of the open space, respectively).

In contrast, for Symmetric triads there was no difference in the probability of jumping first of jumpers in the *diagonal* (16.1%) and *behind* (10.4%) hoop positions, with the probability of the jumper *in front of* the open space was much more likely to jump first (73.5%). That is, Bonferroni post-hoc analysis revealed that a jumper *in front of* the open space jumped first significantly more (both p*<*.05) than the jumpers in the other two positions.

As for the triangle workspace, the results for the square workspace for symmetric triads are consistent with both W*∩*JT and W*∩*JP in Table 1 for a JT and JP of *S*_3_ and a workspace symmetry of *D*_4_. That is, for symmetric groups the same pattern of results observed in [12] were observed, with the coordination pattern reflecting *D*_1_(*Z*_2_), which is the highest order isotropy subgroup of *D*_4_*∩S*_3_. On the other hand, the results for the asymmetric triads, at first glance, appeared to align with the predictions of W*∩*JT (*I*) rather than W*∩*JP (*D*_1_(*Z*_2_)), suggesting that trait symmetry shaped the observed behavioral pattern. Although the classroom leader did not exclusively decide jumping direction, the presence of this leader still operated as a symmetry breaking factor, above and beyond that introduced by *S*_3_ in *D*_4_.

### 2.3. Triads’ synchrony revealed by temporal measure of triadic jumping

The mean difference in the temporal lag are shown in Fig. 3A. (i.e., the average of the temporal lags rom the time the first child jumped to the time the second and third children jumped, respectively). This temporal lag was analyzed using a two-factor mixed-design analysis of variance, with Triad-symmetry (symmetric vs asymmetric) as a between-triad factor and geometrical symmetry of workspace (Geo-symmetry: triangle vs. square) as within-triad factor. The analysis revealed no significant effect of Triad-symmetry (F(1,20)=0.061, p=0.807, 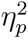=0.003, 1-*β*=0.064) and a significant effect of the Geo-symmetry of workspace (F(1,20)=14.707, p=0.001, 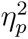=0.424, 1-*β*=0.999). An interaction between Triad-symmetry and Geo-symmetry was also significant (F(1,20)=12.045, p=0.002, 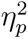=0.376, 1-*β*=0.998). A simple main effects analysis revealed a significant effect of Geo-symmetry of workspace for symmetric triads (F(1, 20)=26.685, p<0.001, 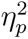=0.572), whereas the effect for asymmetric triads was not significant (F(1,20)=0.066, p=0.799, 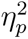=0.003).

**Figure 3:**
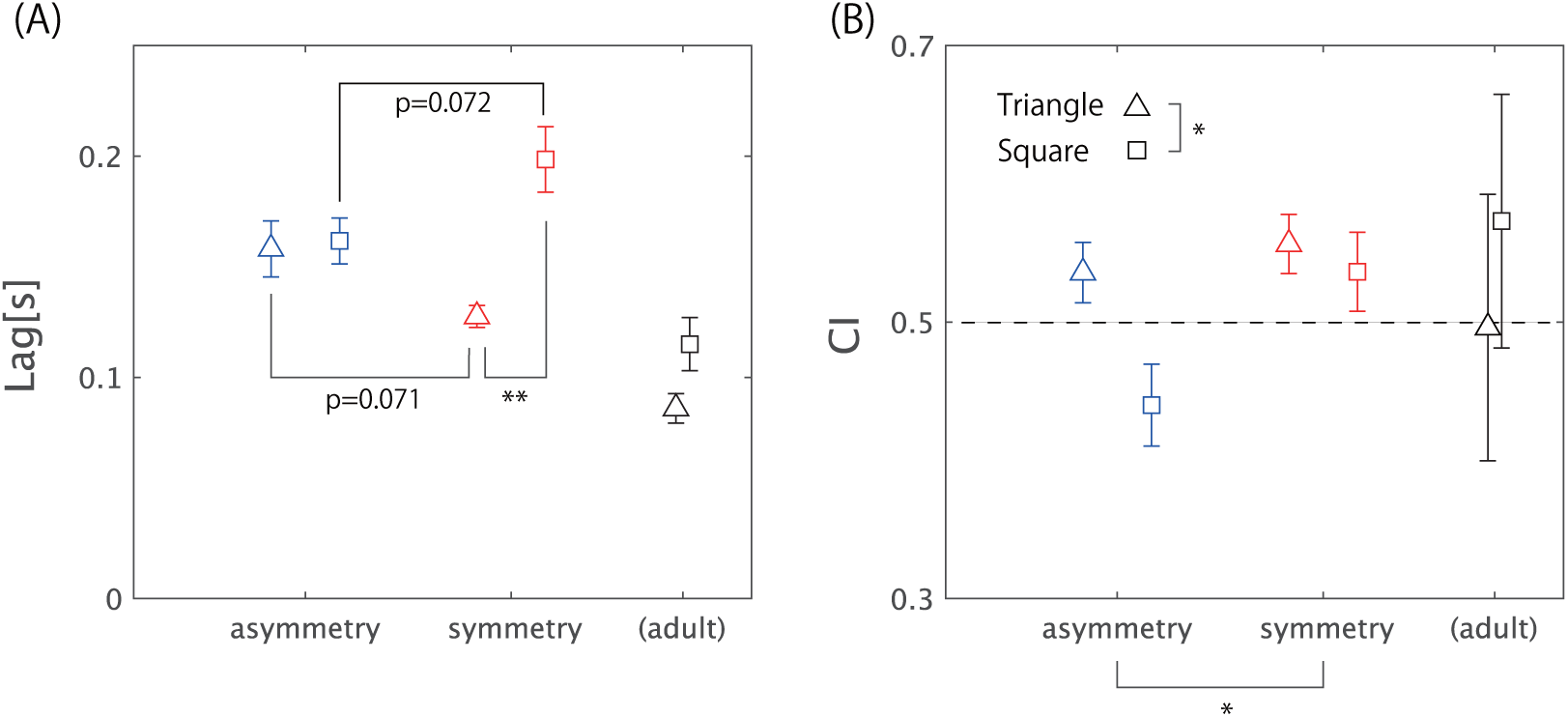
Temporal performance of jumping coordination. **(A)** Lag in triads’ synchrony and **(B)** coordination index for asymmetric and symmetric child triads and for the adult triads measured in previous research[12], as a function of the Triangle and Square workspaces. *: p*<*0.05, **: p*<*0.01

As shown in Fig. 3(A), a difference in temporal lag was observed between the triangle and square workspaces for symmetric triads only. This trend resembles that observed in adults, which is attributed to differences in the symmetry of the workspace-jumper configuration. The relatively shorter delay in the triangle workspace may be due to the symmetry of the jumper positions, as each jumper had no open space on either side. Conversely, the relatively long delay in the square workspace could be attributed to a lead jumper positioned *in front of* an open space, who jumped significantly earlier than the others. In contrast, for asymmetric triads, classroom leaders, who were likely to act as lead jumpers, tended to initiate the jump significantly earlier than the other two in the triangle workspace. Furthermore, in the square workspace, these leaders appeared to shorten the lag between themselves and the lead jumper. However, in both workspaces, the simple main effect of triad symmetry on jumping lag was slightly below the significant level (triangle: F(1,20)=5.100, p=0.071, 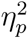=0.203, square: F(1,20)=4.184, p=0.072, 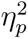=0.173).

As an additional measure of temporal coordination, we also examined the ratio of the lag between the lead jumper and the 2nd jumper to that between the lead jumper and the 3rd jumper. This *coordination index; CI* measure is shown in Fig.3(B), and indicates the temporal coupling between three jumpers. In short, a value closer to 0 indicates that only third jumper jumps much later compared to the first and second jumpers, while a value close to 1.0 indicates that the second and the third jumper jumped almost simultaneously (i.e., at the same lag form the first jumper). A value close to 0.5 indicates that the three jumpers are almost equally spaced in time (i.e., the time lag between the second and third jumpers was the same as the lag between the first and second jumpers).

As can be seen from an inspection of Fig. 3(B), adult jumpers in [12], exhibited CI values around 0.5 in the triangle workspace and slightly above 0.5 in the square workspace. For the children examined in this study, the CI in the triangle workspace was again around 0.5, indicating that the temporal lag between each jumper was (more or less) symmetric (the children jumped 120 degrees out of phase). In contrast to the adult data, however, the CI for the asymmetrical child jumpers was below 0.5, most notable in the square condition. Indeed, an analysis of the CI using a two-factor ANOVA revealed a significant main effect of Triad-symmetry (F(1,20)=5.424, p=0.031, 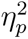=0.213, 1-*β*=0.916), with CI for the asymmetric triads (0.488) significantly smaller than that for the symmetric triads (0.546). The main effect of Geo-symmetry was also significant (F(1,20)=4.904, p=0.039, 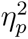=0.197, 1-*β*=0.889), with CI for the square workspace (0.488) significantly smaller than that for the triangle workspace (0.546). The interaction was not significant (F(1,20)=2.117, p=0.161, 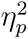=0.096, 1-*β*=0.552).

A close examination of the data indicated that the coordination lag of the child jumpers tended to be greater than that of the adults. However, like the adult triads, the children in the symmetric triads did not *react* to the direction of the leader’s jump, but *synchronized* as close as possible their preparatory movements with the lead jumper to “propel” the triad in the direction of the jump. Evidence of this was provided by the fact that the average delay time of symmetric triads was extremely short, less than 0.2 seconds (the time from the signal that triggers the jump to the time the foot takes off)[25], which is even shorter than the choice reaction time of finger movements; i.e., *≈* 0.2 to 0.3 seconds[26]. Furthermore, the trajectories of the three jumpers, shown in Fig. 1(A), indicate that they were synchronized from the preparatory phase of takeoff; all three jumpers consistently moved their entire bodies up and down in a Mexican-hat-shaped trajectory.

However, the temporal order of the asymmetric triads differed from that of the adults and symmetric child triads. In particular, the classroom leader was significantly ahead of the other two in the triangle workspace and jumped almost immediately after (or as close to the same time as possible) the leading jumper in the square workspace. This finding aligns with the first-jumper probability results reported above, further indicating that the presence of the classroom leader influenced the observed coordination pattern, even when the workspace’s symmetry constrained the action capabilities of the child designated as a classroom leader.

Additional analyses of spatial performance, including synchronization misses, jumping-direction bias, and the local complexity of jumping-direction sequences, are provided in Appendix B. The effect of school grade on the local complexity of jumping-direction sequences is reported separately in Appendix C.

## 3. Discussion

In this study, we investigated whether symmetry principles could be used to understand the patterns of triadic jumping that emerge when children with distinct leader-follower traits performed a group jumping task in triangular and square workspaces. We hypothesized that symmetrical triads (all members with moderate or equivalent leader-follower tendencies) would exhibit coordination patterns consistent with higher-order isotropy subgroups *D*_3_(*S*_3_) in the triangle workspace and *D*_1_(*Z*_2_) in the square workspace, mirroring previous findings with adult triads [12, 27]. More importantly, however, we significantly extended previous research by also examining whether asymmetries in leadership traits (as assessed by classroom teachers) further modulated the patterns of behavioral coordination that emerged. More specifically, we explored whether these trait asymmetries acted as a symmetry-breaking factor in the coordination patterns that formed among the children, above and beyond the positional or geometric (workspace) (a)symmetries already imposed by the task.

As predicted, the results demonstrated that both workspace geometry (triangular vs. square) and the leader-follower traits of the jumpers jointly determined who jumped first in the triadic jumping task. For *symmetric triads* (all members rated as moderate/equivalent), the observed coordination patterns aligned with the higher-order isotropy subgroups specified in Table 1 for *W ∩ JT* and *W ∩ JP* in the first and second rows merged under the label *Adult & Symmetrical Children*. Specifically, in the triangle workspace *D*_3_(*S*_3_), each participant was equally likely to lead *D*_3_(*S*_3_), whereas in the square workspace *D*_4_, a single leader consistently emerged *D*_1_(*Z*_2_). These outcomes replicate previous adult triad findings and match the predictions of W*∩*JT and W*∩*JP for groups with *S*_3_ trait symmetry.

Conversely, when *asymmetric triads* included a designated classroom leader, trait asymmetries further shaped the group’s behavior accordingly. Importantly, the observed coordination patterns were not fully predicted by JT alone. In the triangle workspace, which had no geometrically unique position, the leader was significantly more likely to jump first, producing a pattern best captured by *D*_1_(*Z*_2_), rather than the *I* symmetry predicted solely from *W ∩ JT* =*I*, and the moderates and followers remained interchangeable corresponding to the *S*_2_ symmetry shown in the JP in bottom two rows merged under *Asymmetrical Children* in Table 1. By contrast, in the square workspace — where the leader was placed in a hoop without adjacent open spaces, while moderates/followers had open spaces on both sides — the resulting pattern resembled the identity subgroup *I*, with this shift indicating that the interplay of both trait asymmetry and positional constraints. Importantly, although the observed subgroup resembled *I*, it did not represent the *I* predicted solely from jumper-trait symmetry (JT). Instead, it reflected an identity-like coordination pattern that emerged from the interaction between trait asymmetry and geometry-defined positional constraints.

Overall, these results demonstrate that both physical (i.e., geometric workspace) and non-physical (i.e., dispositional traits) constraints operate as symmetry-breaking factors, but in a hierarchical manner: environmental geometry first constrains the available coordination modes, whereas leader-follower dispositions subsequently bias the selection among those modes. By situating these empirical findings within the framework of symmetry, our study underscores the broad applicability of symmetry and symmetry-breaking principles for understanding the emergence of coordinated actions within small groups.

As noted in the introduction, some form of symmetry breaking was essential for effective task performance in the triadic jumping task employed here. Indeed, symmetry breaking is fundamental to the effective initiation of functional movement patterns. This has been demonstrated extensively in individual animal locomotion [17, 28], wherein symmetry breaking transitions often depend on energetic [29], kinetic, or morphological constraints [30, 31]. For instance, in most quadruped gaits, the legs move out of phase so that at least one leg is always in the air while another leg remains on the ground (except in “pronking,” where all four legs take off and land together) [15, 17]. This asymmetry enables forward propulsion, stability, and acceleration by retaining a critical degree of freedom for movement initiation, even though all legs may momentarily be airborne at higher speeds.

Similar to quadruped gait systems, triadic jumping initiates movement through a breaking of symmetry in the geometry–jumper trait configuration. In the square workspace, the open space introduces a critical degree of freedom that effectively designates a leader; geometric constraints dominate over individual dispositions in determining who jumps first. In contrast, the triangle workspace provides no such open space, so geometry alone does not establish a leader. Instead, the stronger leadership disposition of the classroom leaders guides the jumping order, while moderates and followers play interchangeable roles, consistent with *D*_1_(*Z*_2_). As shown in the third row in Table 1, this outcome matches the symmetry-based predictions for the triangular condition, underscoring how the interplay of geometry and trait asymmetries governs the initiation and maintenance of triadic jumping behavior.

Symmetry-based reasoning also applies to coordination within the human motor system more generally [32, 33]. In bimanual coordination, for instance, each limb has two degrees of freedom (flexion and extension), and under high temporal demands, an in-phase pattern often emerges as the system’s highest-order symmetry. Likewise, similar principles appear to govern cooperative behavior between individuals connected by optical information [5, 6], as modeled using systems of partial differential equations [34, 35].

While our findings revealed small differences from adult triads in the temporal order of the children’s jumps (Fig. 3; see also Supporting Information Text 2), elementary school students (aged 6-10 years) were nonetheless fully capable of completing the triadic jumping task successfully. This observation aligns with prior research indicating that even preschool-aged children can engage in joint actions [36, 37] and that infants as young as 18 months can already recognize others’ goals [38, 39, 40, 41, 42, 43]. By around three years of age, children begin to understand intentions that differ from their own [44], can compensate for others’ actions [45], and even regulate roles to achieve shared outcomes [46]. Critically, these cooperative capacities develop most robustly in human–human interactions, rather than when coordinating with robots [47, 48] or solely rhythmic audio cues [37], and they reflect the maturation of neural mechanisms for motor inhibition [49], mentalizing [50, 51, 52, 53, 54], and action perception [55, 56, 57].

Our data suggest, however, that flexibility in predicting others’ actions continues to evolve throughout the elementary school years. Indeed, sequences of left-right jumping directions were more complex in fourth graders (about 10 years old) than in second graders (about 7 years old) (Supporting Information Text 3). This observation is consistent with previous findings suggesting that five-year-olds have not yet achieved fully flexible action prediction [58]. Although a detailed exploration of this developmental trajectory lies beyond the scope of this paper, our results underscore the nuanced interplay between emerging cognitive-motor skills and the constraints imposed by both leader-follower traits and workspace geometry.

In conclusion, our findings reveal that external constraints play a pivotal role in determining the coordination solutions adopted by triadic systems, which are inherently more complex than dyadic systems. Compared to dyadic interactions, triadic systems exhibit greater instability due to their wider range of possible coordination patterns [59], and these “network motifs” [60] appear to be shaped more by the geometry of the environment than by the coupling among the three actors. Although this study focused on triadic jumping, the algebraic principles of symmetry (i.e., group theory) outlined here can be extended to more diverse multi-agent scenarios, including realworld crowd dynamics [61, 62] and tasks where individuals must navigate workspaces with physical and informational task constraints, while simultaneously adapting to cooperative or competitive relationships [63, 64, 65]. Indeed, we have already replicated these triadic patterns in virtual environments [27], and are developing systems in which virtual agents adjust their leader-follower tendencies in response to human participants. Thus, the current methodology provides a promising way of disentangling the interplay between environmental constraints and actor-specific traits, offering a powerful means to investigate and predict coordinated behaviors in larger and more complex groups.

### Methodological and theoretical significance

Our group-theoretic framework offers a compact formalism for predicting coordination outcomes across domains. By treating interpersonal dynamics as constrained by discrete symmetry groups, this framework allows a finite set of possible coordination modes to be derived without simulating complex neural or social models. This approach could generalize to other contexts, including team sports, collaborative robotics, and crowd dynamics, where environmental and social asymmetries interact.

### Limitations and future directions

The current study used a simplified task and a limited age range. Future research should examine more diverse participant groups and ecological settings, integrate physiological measures, and test adaptive virtual agents capable of dynamic symmetry manipulation. We also plan to analyze how coupling strength and feedback delay influence emergent leadership patterns, expanding toward multi-agent modeling and neural coupling analyses.

## Supporting information

App

## 4. STAR Methods

### Resource Availability

#### Lead Contact

Further information and requests for resources should be directed to Akifumi Kijima.

#### Materials Availability

This study did not generate new materials.

#### Data and Code Availability

Data and analysis scripts will be made available upon publication via https://github.com/akijima-oss/Data_JumpTriads.git.

### Method Details

#### Participants

Forty-seven children, 23 second-graders and 24 fourth-graders participated in the experiment. All second- and fourth-grade participants were recruited from the University of Yamanashi Elementary School. None of the children had a history of major physical or psychiatric illness, and all children in each grade were recruited from the same classroom within the same school. After obtaining approval from the Research Ethics Committee of the Faculty of Education, University of Yamanashi, informed consent was obtained from the participants and the participants’ parents through the staff of the University of Yamanashi Elementary School.

#### Formation of Asymmetric and Symmetric triads

Participants were assigned to one of two types of triads―Asymmetric triads or Symmetric triads― based on their tendency to take on leader or follower roles in daily life. This tendency was assessed using a custom one-item questionnaire designed to measure each child’s inclination toward leader or follower roles in everyday classroom activities. Each child’s classroom teacher rated them on a scale from −3 (“always behaves as a follower”) to +3 (“always behaves as a leader”), with 0 indicating “does not take a particular role.” Each of the 11 Asymmetric triads consisted of three children from the same grade: one rated as −3 (strong follower), one as +3 (strong leader), and one rated near 0 (neutral). In contrast, each of the 11 Symmetric triads was composed of three children all rated near ± 1 or 0. Based on the results of this assessment, we formed five Symmetric triads and five Asymmetric triads for the 2*^nd^* graders, and six Symmetry and six Asymmetric triads for the 4*^th^* graders.

#### Triadic jumping task

Three or four plastic hoops, each with a diameter of 0.6 meters, were placed at the center of a 2.28 × 2.28 meter polyurethane mat. As illustrated in Fig. 1(A) and (B), the hoops were arranged to form either a triangle or a square, with each hoop touching its adjacent hoops so that the center-to-center distance between adjacent hoops was approximately 0.6 meters. Each member of the triad was assigned to one of three colored hoops ―yellow, blue, or red―and instructed to jump to the left or right hoop with both feet in response to a jump signal presented by an electronic metronome (see Supporting Information Movies S1 and S2).

Two preliminary signals were given at 1.0-second intervals before the jump signal, resulting in a jump action cycle of approximately 3.0 seconds, including a 2.0-second preparation phase, as depicted in Fig. 1(A). This periodic jumping movement continued for a total of 15 jumps. If any participant in a triad collided with another, the attempt was considered unsuccessful, and the participants were instructed to return to their assigned starting hoop and restart the sequence.

Six triads from each of the Asymmetry and Symmetry groups began with the triangle condition, while the remaining five started with the square condition. The workspace geometry was changed after the triads successfully completed 15 jumps in the initial condition. Participants were not provided with any instructions regarding the direction of their jumps. Instead, they were explicitly instructed not to communicate verbally or nonverbally during the experiment and not to assign leader or follower roles before each jump or trial (see, Supporting Information Movies S1 and S2).

Participants were told to freely choose their jumping direction at each jump based on individual decision-making, with the understanding that all members of the triad needed to jump in the same direction to avoid collisions. Consequently, each member had to anticipate the jumping direction of the other two by synchronizing their movements during the 2.0-second preparatory phase, which involved a downward shift of the center of mass and a forward/upward arm swing to generate sufficient ground-reaction impluse for the 0.6-meter jump between hoops. All methods were performed in accordance with the Declaration of Helsinki.

#### Workspace geometry

The aim of the experiment was to test how the workspace and trait (a)symmetries influenced the jumping patterns of the child triads and the degree to which they were consistent with the symmetry group predictions outlined in Tables 1 and Figure 1. In Symmetric triads, three jumpers with comparable moderate leader-follower trait (*S*_3_) were randomly placed into three or four hoops as depicted in the upper panels of Fig. 1(C). Under this alignment, we expected that the results would be consistent with our previous research [12, 27] and that the observed jumping patterns would be consistent with *D*_3_(*S*_3_) for the triangle workspace and *D*_1_(*Z*_2_) for square workspace.

For the Asymmetric triads, the positioning of the triad members was arranged so that the classroom leader was always placed between the children with moderate and follower tendencies (as illustrated in the lower panels of Fig. 1(C)). In the square workspace, this setup ensured that the child designated as the classroom leader was always positioned in a hoop with no open space on either side. That is, the children designated as moderates and followers were placed with equal probability in one of the two hoops that had an adjacent open space (yellow and red in Fig. 1(C)).

This experimental setup allowed us to separately analyze the impact of the asymmetry in leader-follower traits on behavioral order, as well as the interaction between this asymmetry and the geometric symmetry of the workspace. For the triangle condition groups, the asymmetry in leader-follower disposition was expected to explicitly break the symmetry, resulting in the child judged to have the strongest leadership tendencies predominantly jumping first. That is, *D*_1_(*Z*_2_) or *I* should be observed, with *D*_1_(*Z*_2_) indicating that, in the presence of a “leader”, moderates and followers could be considered equivalent (i.e., non-leaders). For the square configuration, where the symmetry was already explicitly broken by the geometric arrangement, two outcomes were possible. First, a jumping pattern consistent with *D*_1_(*Z*_2_) could be observed, indicating that the coordination pattern that emerged was unaffected by the leader-follower asymmetries and simply emerged as a function of W*∩*JP. However, if asymmetries in jumpers’ traits did play a role in defining the pattern of coordination that results (i.e., is defined by W*∩*JT) then a coordination pattern consistent with *I* would be observed (see Table 1 and Supporting Information Text 1, “Detail of Symmetry Calculation”).

Importantly, these different possibilities could be assessed by calculating the probability of each individual jumper becoming the lead jumper, as a function of the variations in leader-follower traits and differences in relative position (see the section below “Each Jumper’s Probability of Being a Leading Jumper”).

### Quantification and Statistical Analysis

#### Data measurement

Eight infrared cameras (Miqus M1, Qualisys, Sweden) were used to record the three-dimensional position of each participant’s head at a sampling frequency of 100 Hz. Each participant wore a plastic helmet with a reflective marker attached to the top of the head. Prior to analysis, the recorded motion data were filtered using a fourth-order Butterworth filter with a cutoff frequency of 6 Hz. To assess the temporal coordination of the triads, each participant’s head height (Z-axis position) was analyzed. The motion data was segmented into individual jump cycles by identifying the peaks (the peak in head positions) over time.

#### Probability of being the lead jumper

The peak head height during each jump for all three jumpers was determined using the time series data depicted in Fig. 1(A). In the first analysis, the probability of being the lead jumper was calculated for each participant according to leader-follower trait category within the Asymmetric triads. Probability distributions were computed separately for the three jumpers in both the triangle and square workspace conditions. A two-factor analysis of variance (ANOVA) was employed to analyze the resulting data, with the geometric symmetry of the workspace (two levels: triangular and square) and jumpers’ traits (three levels: classroom leader, moderate, and follower in Asymmetric triads) as within-triad factors. In the second analysis, the probability of being the lead jumper was calculated for each participant across the different hoop positions for both Asymmetric and Symmetric triads. A two-way mixed-design ANOVA was then employed to analyze the resulting data, with triad symmetry as the between-triad factor and jumper position as the within-triad factor. It is important to note that hoop position is irrelevant in the triangle workspace due to the symmetrical jumper-geometry configuration. However, in the square workspace, the hoop placed diagonally opposite the open space is unique, as it lacks an adjacent open space, distinguishing it from the other two positions. For Symmetric triads, the probability of being the lead jumper is the same (*D*_3_(*S*_3_)) among the three jumpers when they jump in the triangle workspace. However, when they jump in the square workspace, only the probability for the jumper positioned diagonally opposite the open space differs from the other two jumpers (*D*_1_(*Z*_2_)) in the square workspace condition (denoted as *W ∩JP* in Table 1). On the other hand, for Asymmetric triads, the hoop positions of the moderates and followers were counterbalanced. Therefore, only the classroom leader, who occupied the unique position, was expected to differ from those of the other two jumpers (*D*_1_(*Z*_2_)) in both workspaces, as denoted by *W ∩ JP* in Table 1.

#### Temporal and spatial measures of coordination

The temporal lag of each of the two following jumpers relative to the lead jumper was calculated for each successful jump (see Fig. 1(A)). These two lags were then averaged to provide an overall estimate of the temporal coordination lag. In addition, a coupling pattern index (CI) was also calculated using these lags as follows. 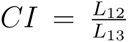 where *L*_12_ = *t*_2_ *− t*_1_ denotes the lag between the first and second jumpers, and *L*_13_ = *t*_3_ *− t*_1_ denotes the lag between the first and third jumpers A CI ratio equal or close to 0.0 indicates a coupling pattern in which the first two jumpers move nearly simultaneously and precede the third jumper, whereas a CI ratio equal or close to 1.0 indicates a coupling pattern in which the second and third jumpers move nearly simultaneously after the first jumper. These temporal coordination metrics were compared using two-way ANOVAs as a function of triad symmetry (asymmetric and symmetric) and workspace symmetry (triangle and square). For the procedures employed for calculating coordination failures and the other spatial measures, see Supporting Information.

### Key Resources Table

**Table 2:** Key Resources Table summarizing instruments, software, and data repositories.

| Reagent or Resource | Source | Identifier |
| --- | --- | --- |
| Motion capture system | Qualisys, Sweden | Miquis M1 and Qualisys Track Manager 2022 |
| Statistical software | R Foundation for Statistical Computing | R 4.3.2 GUI 1.80 Big Sur Intel build (8281) |
| Dependent measure calculation scripts | Mathworks Inc., USA | MATLAB R2022b (maci64) |

## Acknowledgments

We thank Keiko Yokoyama and Keisuke Fujii for their valuable comments on the experimental design, and Satoshi Suzuki for his advice and assistance with coordinating the children’s participation. This work was supported by JSPS KAKENHI Grants 20H04090, 22K19727, 23K20369, and 25H01104.

## Author Contributions

A.K., M.O., H.S., Y.Y., R.W.K., and M.J.R. designed the research. A.K., M.O., and Y.Y. performed the experiments. A.K., M.O., H.S., Y.Y., R.W.K., and M.J.R. analyzed the data. A.K., H.S., R.W.K., and M.J.R. developed the symmetry analyses. A.K., R.W.K., and M.J.R. wrote the manuscript. R.W.K. and M.J.R. contributed to the interpretation of the results and critically revised the manuscript.

