## Supplementary material for "Symmetry breaking and social coordination in children: Using group theory to understand emergent patterns of multi-agent coordination": App

Akifumi Kijima et al.

**Corresponding author:** Akifumi Kijima
Faculty of Education, University of Yamanashi, Japan


#### Appendix A. Detail of symmetry group calculation

Based on group theory concerning symmetry, we predict a cooperative order where leader-follower dispositional traits exhibit differences between symmetrical and asymmetrical triads within both triangular and square workspaces. Fig. A.1(A) illustrates the two workspace dihedral symmetries. Dihedral symmetry refers to the set of all isometries — angle- and distance-preserving rigid motions, namely rotations and reflections — that leave a regular polygon invariant. For a regular $n$-gon, the associated dihedral group $D_n$ consists of $2n$ such transformations: $n$ rotations by integer multiples of $2\pi/n$ and $n$ reflections about its symmetry axes. In other words, dihedral symmetry characterizes the full geometric invariance of a regular polygon under both rotational and reflectional isometries. The equilateral triangle workspace, constructed by placing hoops of equal diameter in mutual contact, can maintain its original shape regardless of whether reflected about any of the three axes in $\mathrm{Ref}_{1-3}$, or rotated by any angle $\mathrm{Rot}_{0,360}$, $\mathrm{Rot}_{120}$, or $\mathrm{Rot}_{240}$. It can therefore be defined as possessing $D_3$ symmetry of order 6. Next, the square retains its original shape when reflected about any of the four axes in $\mathrm{Ref}_{1-4}$, or rotated by any angle from $\mathrm{Rot}_{0,360}$, $\mathrm{Rot}_{90}$, $\mathrm{Rot}_{180}$, or $\mathrm{Rot}_{270}$. It can thus be defined as an order-8 $D_4$ symmetry.

Fig. A.1(B) illustrates the jumpers' trait symmetry and jumpers' position symmetry for Symmetrical triads and Asymmetrical triads respectively. These symmetric groups on $n(=3$ in this experiment) elements, denoted $S_n$, are the permutation group of all bijections of an $n$-element set onto itself. In other words, each element of $S_n$ is a permutation — a bijective map that reorders the $n$ elements. The order of the group is $n!$, because there are $n!$ possible permutations of $n$ distinct elements. Both symmetries for Symmetrical triads in the upper row are identical and correspond to $S_3$ with six elements ($I$ and $\mathrm{P}_{1-5}$), however, those symmetries for Asymmetrical triads differ. For Asymmetrical triads, each jumper's dispositional trait itself was different from the others and cannot be permuted at all, so jumper trait symmetry is $I$. However, jumper-position symmetry is defined by the arrangement of the jumpers across the hoops. As demonstrated in the text, due to the counterbalancing of Moderates and follower positions, the Jumpers' position symmetry in Asymmetrical triads differs from jumper-trait symmetry (JT) and is $S_2$ with only two elements including $I$ and $\mathrm{P}_1$.

Fig. A.1(C) depicts the highest-order dihedral group within the common subgroup of the workspace symmetry and the jumpers ' positional (JP) symmetry. For a triangle workspace where the number of jumpers equals the number of hoops, all three hoops have no adjacent open space and therefore share equal degrees of freedom. If the JP symmetry is itself symmetric, the three jumper-hoop configurations form the dihedral group $D_3$, as shown in the upper-left panel. If the JP symmetry is asymmetric ($S_2$), the configuration reduces to $D_1$ of order 2, as shown in the lower-left panel. Conversely, when the workspace is square, only two of the three hoops (red and yellow) possess adjacent open spaces. As a result, the degrees of freedom for the triad correspond to $D_1$ of order 2. This applies to both symmetrical and asymmetrical triads. In these triads, because the red or yellow hoops were assigned to moderates and followers with equal probability, the JP symmetry is $S_2$. Nevertheless, the maximal-order subgroup common to the workspace symmetry $D_4$ and the jumper-position symmetry $S_2$ is $D_1$ (isomorphic to $Z_2$).

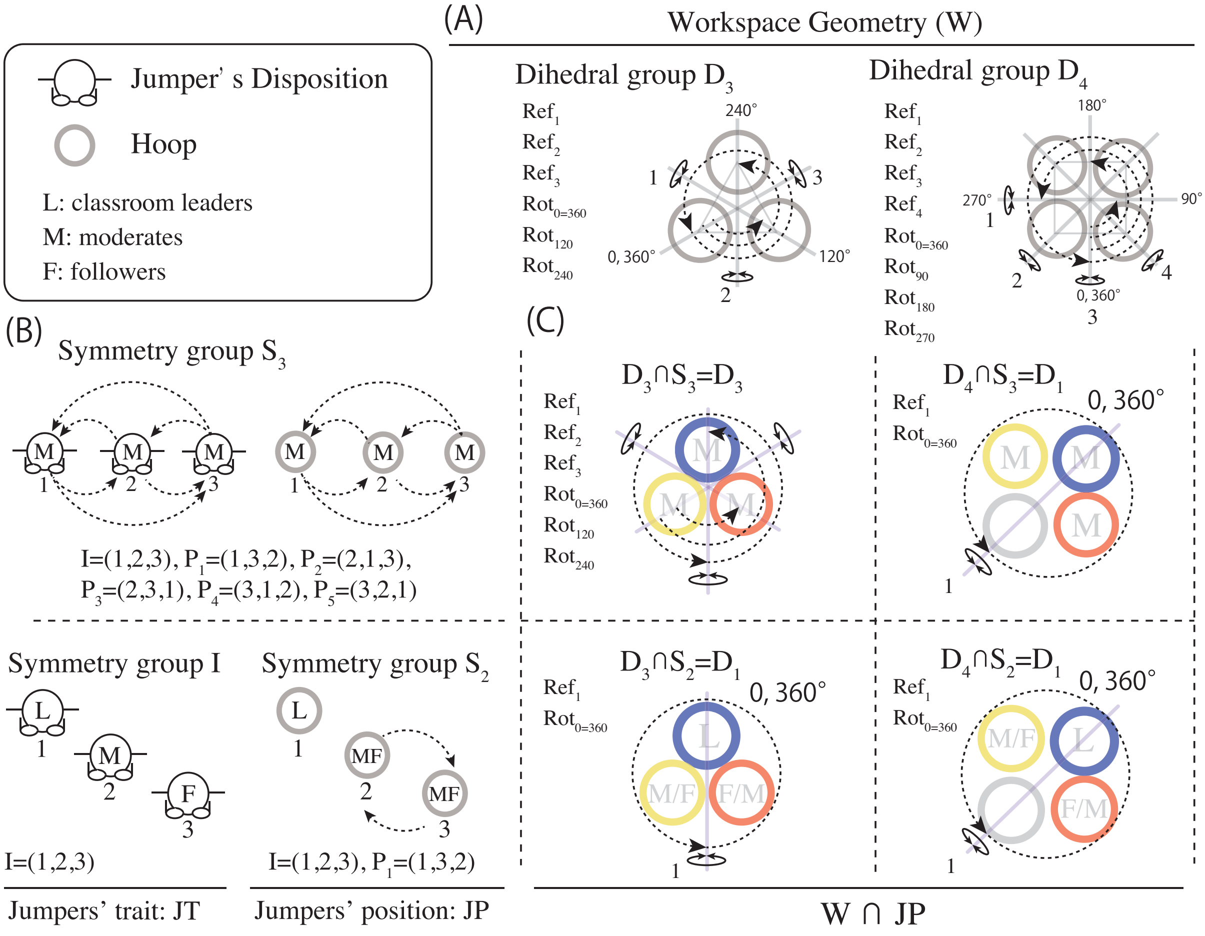


Figure A.1: (A) Dihedral group ($D_n$) of the workspace geometry. (B) Symmetry group ($S_n$) of the jumpers' trait symmetry and jumpers' position symmetry. (C) Jumper-hoop configuration illustrating, for each dispositional leader-follower trait, the highest-order common subgroup of the workspace symmetry and the jumpers' positional symmetry (W ∩ JP).

### Appendix B. Spatial performance of triadic jumping

*Methods*

To assess spatial performance, we quantified the number of "synchronization misses" by counting the total collisions that occurred as each triad attempted to complete 15 successful jumps. This metric served as an index of triad synchrony. We also recorded the percentage of counterclockwise (i.e., rightward) jumps among the 15 successful jumps.

Additionally, we measured the local complexity[1, 2] of each sequence of 15 jumps as an approximation of its algorithmic complexity, using the coding theorem method[3]. The method relies on the relation ($K(s) \approx -\log_2(D(s))$), where $K(s)$ represents the Kolmogorov (algorithmic) complexity of a string (s), and $D(s)$ denotes its algorithmic probability. To estimate $D(s)$

for each sequence of 15 left-right jump directions (e.g., `RRRLRLRLRLRLRRL`), we employed a large-scale simulation of Turing machines (approximately 4.5 million) using the `.acss` package in the R statistical programming language (The R Foundation for Statistical Computing). Each Turing machine was simulated until it either halted or did not halt. Since the .acss package supports a maximum sequence length of 9, we divided each 15-direction sequence into seven overlapping subsequences of length 9. We then calculated the local complexity for each subsequence and averaged those values to obtain the overall local complexity score for each triad ' s jump-direction sequence.

*Results*

As shown in Fig. B.2(A), the analysis of synchronization misses revealed no main effects of (Triad-Symmetry: F(1,20)=0, p=1, $\eta_p^2$=0, 1-$\beta$=0.050; Geo-Symmetry: F(1,20)=0.526, p=0.476, $\eta_p^2$=0.026, 1-$\beta$=0.429), nor an interaction between Triad-Symmetry and Geo-Symmetry (F(2,26) = 0.999, p = 0.329, $\eta_p^2$=0.048, 1-$\beta$=0.486). That is, the success rate of triadic jumping was not influenced by the geometric symmetry of the workspace, nor the leader-follower symmetry within the triads. It should be noted, however, that the success rate of the child triads in the current experiment was higher than that observed for the adult triads reported in [4, 5]. This difference in performance by the child triads could be attributed to the greater number of practice sessions provided for them to learn to synchronize their jumps; adult triads were only provided 2-3 practice sessions, whereas child triads were allowed until they were able to synchronize their jumps successfully.

Fig. B.2(B) shows the results for the percentage of counterclockwise jumps. Again, there was no main effect of Triad-Symmetry (F(1,20)=0.284, p=0.600, $\eta_p^2$=0.014, 1-$\beta$=0.176) nor Geo-Symmetry (F(1,20)=0.003, p=0.958, $\eta_p^2$=0, 1-$\beta$=0.005), nor was there an interaction effect (F(2,20)=0.999, p=0.329, $\eta_p^2$=0.048, 1-$\beta$=0.486). Finally, Fig. B.2(C) shows the results for K9 complexity, where there was also no main effect of Triad-Symmetry (F(1,20)=0.005, p=0.945, $\eta_p^2$=0, 1-$\beta$=0.052), or Geo-Symmetry (F(1,20)=0.248, p=0.624, $\eta_p^2$=0.012, 1-$\beta$=0.233), nor was there an interaction effect (F(2,26)=0.003, p=0.958, $\eta_p^2$=0, 1-$\beta$=0.051).

Collectively then, these results indicated that there was no effect of experimental condition of triads and that jumping direction was not influenced by the dispositional/trait symmetry of triads, nor the geometric symmetry of the workspace. Moreover, there was no significant difference in spatial bias between child and adult triads, as both groups exhibited a strong preference for the counterclockwise direction, with 70% of successful coordinated jumps occurring in that direction. The decision on which direction to jump seemed to arise spontaneously—indicative of a spontaneous symmetry breaking—during each jump. While the nature of this synchrony is discussed in the main text, it still does not fully account for the overall shared preference for counterclockwise jumping over clockwise.

### Appendix C. Effect of school grade on the local complexity of jumping direction sequences

Fig. C.3 presents the complexity of fifteen sequences of jumping direction as a function of grade (2nd vs 4th), triad symmetry and workspace geometry. A three-way analysis of variance (ANOVA) was conducted with Triad-Symmetry and Grade (2nd and 4th) as between-participants factors and Geo-Symmetry as a within-participants factor. The results indicated that the main effect of Triad-Symmetry was not significant ($F(1,18) = 0.024$, $p = 0.879$, $\eta_p^2$=0.001, $1-\beta = 0.053$), nor was the main effect of Geo-Symmetry ($F(1,18) = 0.178$, $p = 0.677$, $\eta_p^2$=0.010, $1-\beta = 0.195$). However, the main effect of Grade was significant ($F(1,18) = 7.468$, $p = 0.013$, $\eta_p^2$=0.293, $1-\beta = 0.845$), with 4th grades (2.71) producing more complex behavioral sequences than 2nd graders (0.79).

Finally, note that there were no interaction effects (Triad-Symmetry x Grade, $F(1,18) = 0.670$, $p = 0.423$, $\eta_p^2$=0.036, $1-\beta = 0.145$; Triad-Symmetry x Geo-Symmetry, $F(1,18) = 0.01$,

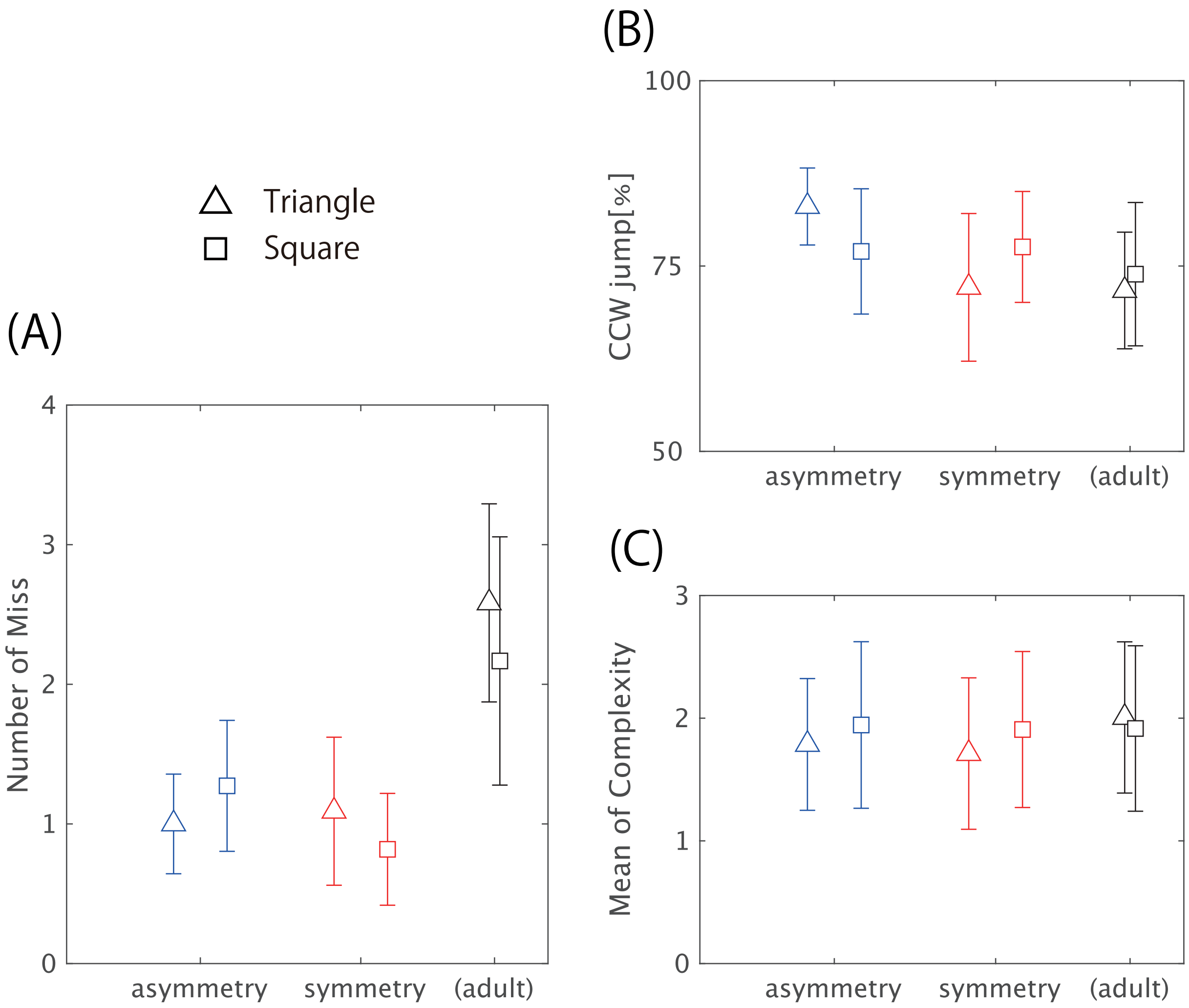


Figure B.2: Spatial performance of triadic jumping. (A) Number of synchronization misses, (B) percentage of counterclockwise jumps, and (C) mean local K9 complexity calculated across the seven overlapping subsequences.

$p = 0.922$, $\eta_p^2$=0, $1-\beta = 0.056$; Grade x Geo-Symmetry, $F(1,18) = 0.385$, $p = 0.542$, $\eta_p^2 = 0.021$, $1-\beta = 0.335$; and Triad-Symmetry x Grade x Geo-Symmetry, $F(1,18)$=0.282, $p$=0.602, $\eta_p^2$=0.015, $1-\beta = 0.259$). These results suggest that differences in grade level influence the complexity of the jumping direction sequence, while the factors of Triad-Symmetry and Geo-Symmetry do not have a significant impact.

[4] A. Kijima, H. Shima, M. Okumura, Y. Yamamoto, M. J. Richardson, Effects of agent-

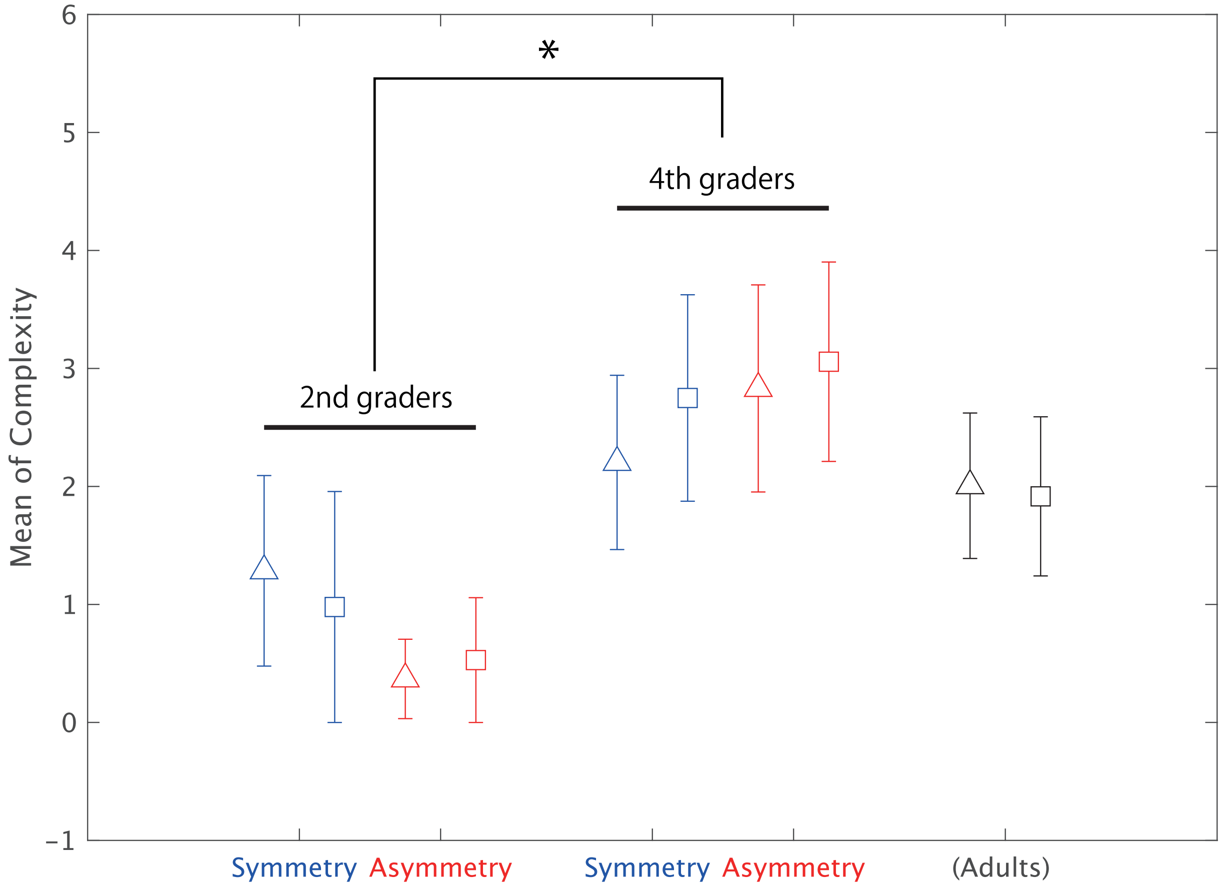


Figure C.3: Difference in K9 complexity related to jumping direction between 2nd graders and 4th graders. *: p=0.01.

environment symmetry on the coordination dynamics of triadic jumping, Frontiers in Psychology 8 (2017) 233637.

[5] A. Naito, K. Go, H. Shima, A. Kijima, Synchrony in triadic jumping performance under the constraints of virtual reality, Scientific Reports 12 (1) (2022) 12417.
